# The functional morphology of hawkmoth lamina monopolar cells reveals mechanisms of spatial processing in insect motion vision

**DOI:** 10.1101/2025.06.19.660550

**Authors:** Ronja Bigge, Kentaro Arikawa, Anna Stöckl

**Affiliations:** Department of Biology, University of Konstanz, Universitätsstr. 10, 78464 Konstanz, Germany; Research Center for Integrative Evolutionary Science, SOKENDAI, Hayama, Kanagawa 240-0193, Japan

## Abstract

Many animals strongly rely on their sense of vision, as it provides information about the natural world with particularly high dimensionality. In insects, the first visual processing stage of the brain, the lamina, plays an important role in parallel processing of this complex information. Its main relay neurons, lamina monopolar cells (LMCs), receive information directly from the photoreceptors and shape the contrast, luminance, spatial and temporal tuning of the insect visual system in a cell-type specific manner. One of their best-investigated downstream targets is the motion vision pathway. However, how LMC types that feed into motion processing delineate contrast and luminance is only known from fruit flies, while the contribution of LMCs to spatial processing has only been described in hawkmoths. Here, we provide a novel characterization of hawkmoths lamina monopolar cells, to integrate the contrast, luminance and spatial processing properties of LMCs in the motion pathway. We used serial block-face scanning electron microscopy to reconstruct the anatomical fine structure of LMCs in a focal lamina cartridge, including their pre- and post-synaptic connections. Combining our novel LMC classification with intracellular recordings, we further investigated the functional role of the main relay neurons to the motion pathway, L1 and L2, in terms of contrast and spatial processing. We show that unlike in flies, L1 and L2 process contrast, and spatial information similarly. Crucially, their two distinct spatial processing functions, lateral inhibition and spatial summation are explained by the density and distribution of their synapses in different lamina layers. Based on these findings, we propose a novel mechanism of delineating distinct spatial processing functions in a single cell.

## Introduction

The visual system acquires sensory information about the environment with particularly high dimensionality, providing luminance, contrast, colour, polarisation information, as well as multiple derived qualities, such as motion or depth information (Land & Nilsson, 2012). To extract relevant features for specific behavioural tasks, neural systems filter and categorize the visual input early on, to reduce its complexity (Niven & Laughlin, 2008). A key feature of such early processing are parallel pathways, to process features simultaneously (Macpherson et al., 2021). In vertebrates, for instance, retinal ganglion cells initialize parallel channels in the retina projecting ultimately to the visual cortex, where spatially defined, cell-specific connectivity recombines these inputs into new parallel outputs (Watanabe & Rodieck, 1989), reviewed in Nassi and Callaway (2009). Despite the structural differences in organization, the insect visual system employs very similar parallel processing strategies (Clark & Demb, 2016).

A prime area in the insect brain where parallel visual processing manifests both morphologically and physiologically is the lamina. As the first processing stage of visual information, it is retinotopically organized into repeated sets of neurons, the so-called cartridges. Each cartridge processes information from one functional “pixel” of the image, typically receiving information from the photoreceptor axons of a single ommatidium in the eye (with the exception of dipteran flies (Braitenberg, 1967; Kirschfeld, 1967)). Depending on species (Strausfeld, 2012), it contains 4-5 types of lamina monopolar cells (LMCs) each, of which processing the photoreceptor input. In forwarding this information to the next brain area, the medulla, they initiate of distinct visual pathways (Cajal & Sanchez, 1915; Ryu et al., 2022; Strausfeld, 1970; Strausfeld & Blest, 1970).

A key pathway that lamina neurons provide input to is motion vision (Borst, 2014). In fruit flies, the two largest LMCs in each cartridge, L1 and L2, initiate two parallel pathways with opposing contrast information: brightness increments and decrements, so-called ON (L1) and OFF (L2) components of the visual signal (Joesch et al., 2010). L1 and L2 are the primary postsynaptic partners of photoreceptors (Rivera-Alba et al., 2011), and have the largest axon diameters and similar branching patterns in the lamina proper (Fischbach & Dittrich, 1989), but project to different layers in the medulla (Meinertzhagen & O’Neil, 1991). Physiological investigations have shown that L1 and L2 have different contrast and luminance coding properties in fruit flies, where L1 encodes a larger luminance component of the visual signal, while L2 encodes primarily contrast (Ketkar et al., 2022; Ketkar et al., 2023).

In addition to luminance and contrast processing, the insect lamina has been implicated in spatial processing since the first descriptions of lamina monopolar cells (Cajal & Sanchez, 1915), because of a prominent morphological characteristic: LMC types in many insect species extend lateral processes into neighbouring cartridges, and are thus ideally suited to either spatially inhibit or pool information from neighbouring visual units. As these processes extend much further in most nocturnal insects than in diurnal ones (Greiner et al., 2004a; Ribi, 1977; Stöckl et al., 2016b; Strausfeld & Blest, 1970; Wolburg-Buchholz, 1979), they were hypothesized to spatially integrate visual information, thereby enhancing the visual signal and reducing associated noise in dim light (Greiner et al., 2005; Greiner et al., 2004b; Meinertzhagen et al., 1983; Strausfeld & Blest, 1970; Strausfeld & Braitenberg, 1970; Warrant, 1999). Since the main relay neurons of photoreceptor information in the fruit fly lamina, L1-L3, lack processes that extend into neighbouring cartridges (Fischbach & Dittrich, 1989; Rivera-Alba et al., 2011), the spatial integration function of LMCs has been mainly investigated in other insects. Evidence for spatial summation was physiologically demonstrated in the so-called type II neurons in the lepidopteran hummingbird hawkmoth, which integrate information from their long processes in dim light, while mainly receiving input from short processes in bright conditions (Stöckl et al., 2020). This dynamic adjustment of spatial tuning and visual sensitivity is measurable in the responses of motion neurons, suggesting that the investigated LMC types in hawkmoths contribute significantly to the motion vision pathway (Stöckl et al., 2017).

However, it is unclear how the spatial processing functions described in hawkmoths align with the contrast and luminance processing contributing to motion vision in fruit flies, since homology between lamina monopolar cells across these species has not been established. To reconcile the contribution of LMCs to spatial, contrast and luminance processing in motion vision across insects, we investigated the anatomical fine structure of all LMCs and photoreceptors in a cartridge of the hummingbird hawkmoth down to the synaptic level. We thereby established a novel classification of LMCs in hawkmoths, proposing homology to other insects with lamina connectome data. We further characterized the contrast and luminance coding properties of individual lamina neurons using intracellular recordings. Finally, we investigated the LMC’s spatial response properties physiologically, and propose how these arise from the LMC’s synaptic connectivity profiles.

## Methods

### Anatomy

#### Tissue preparation and image acquisition

The tissue preparation for serial block-face scanning electron microscopy (SBF-SEM) was performed as previously described in (Deerinck, 2010) and will briefly be summarized here. We dissected the brains out of the head capsule and pre-fixated them in 4% glutaraldehyde and 1% paraformaldehyde in 2mM CaCl_2_ in 0.15M sodium cacodylate buffer (pH 7.3, Ca-CB). The dissection and pre-fixation was performed at Würzburg University, Germany, and the following procedures at Sokendai, Japan. The brains were then washed in Ca-CB, embedded in 35% gelatine-filled mold and cut into 150 µm thick sections using a vibrating microtome (VT-100S, Leica). The sections were stored in Ca-CB overnight at 4 °C. The next day, the sections were treated with 2 % osmium tetroxide (OsO_4_) and 1.5 % K-ferrocyanide in Ca-CB for 1 h on ice. Then, they were washed with Milli-Q water (MQ, 5 x 3 minutes at room temperature). Subsequently, the sections were treated with 1 % thiocarbohydrazide in MQ (20 minutes at room temperature) and then, washed with MQ (5 x 3 minutes at room temperature). After that, the sections were transferred to 2 % OsO_4_ in MQ for 30 min at room temperature. They were then washed with MQ (5x 3 min at RT) and subsequently transferred to 1 % uranyl acetate in MQ at 4 °C overnight. The next day, the sections were transferred to Walton’s lead aspartate solution for 30 min at 60 °C. Finally, the sections were then dehydrated in an increasing ethanol series (5 min each on ice), treated with acetone, and embedded in Spurr’s resin (Polyscience, Inc.).

The serial images were obtained using SBF-SEM (Zeiss Gemini SEM300, Gatan 3 View). The samples were aligned such that the images were taken perpendicular to the cell axon’s longitudinal axis. In total, about 3000 images were collected at a 70 nm interval in z-direction, covering 80 x 80 µm in x-y direction (10,000 × 10,000 pixels, pixel size = 8 nm). The images contain about 70 cartridges in total and cover the cells from the cell body layer, along the full extent of the lamina to the first optic chiasm (Fig 1A).

**Figure 1:**
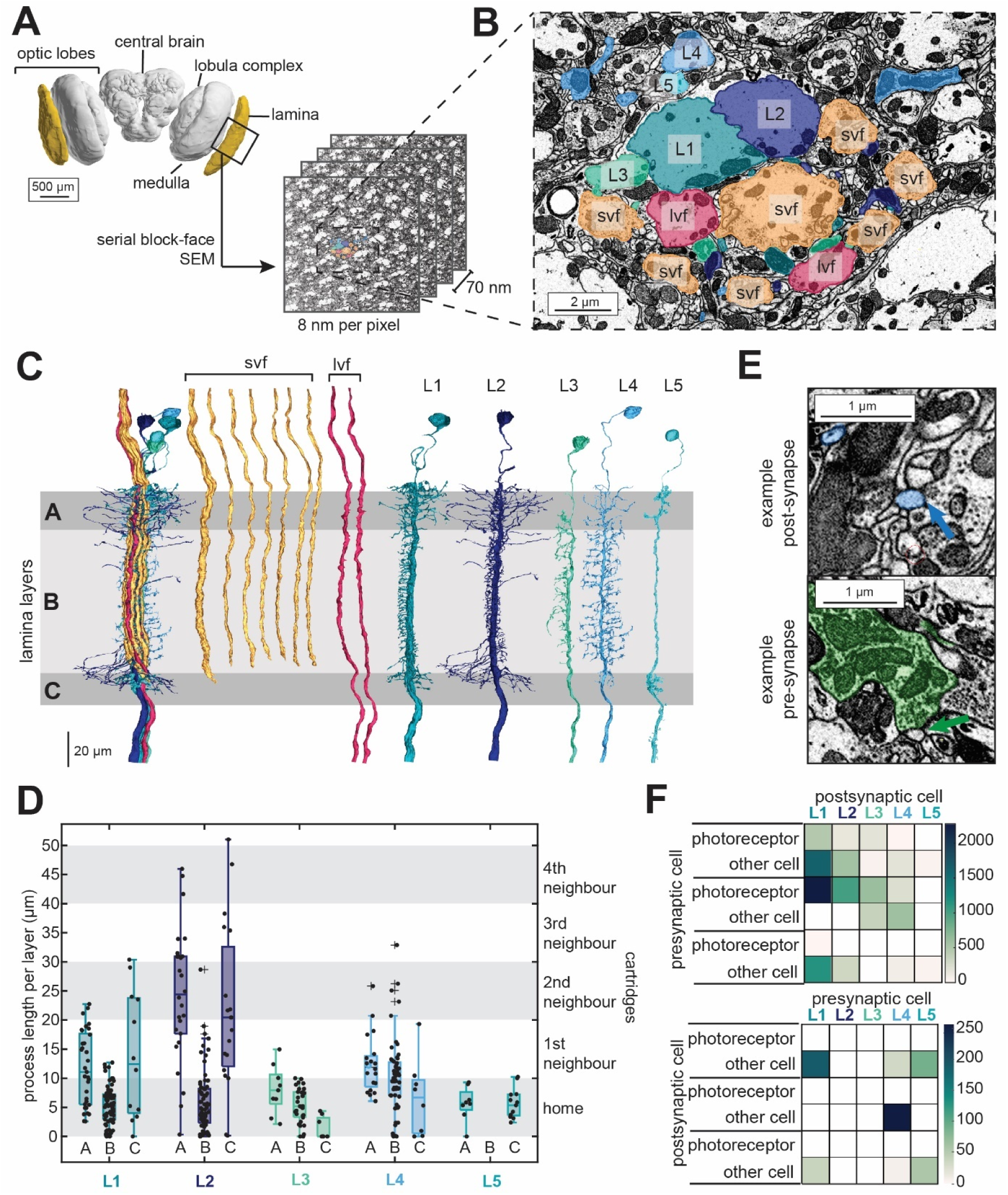
Anatomical reconstruction of a lamina cartridge in the hummingbird hawkmoth. **A** 3D volume rendering of the hummingbird hawkmoth brain; the lamina is highlighted in yellow. The position of the lamina section analysed is approximated by the black box. The image stack obtained by serial block-face electron microscopy consisted of 3018 images of 10,000 × 10,000 pixels, with a pixel size of 8 nm. The intersection distance between the images was 70 nm. **B** Zoomed-in cross-section of the lamina highlighting the focal cartridge that was 3D reconstructed and analysed (in colour). **C** 3D reconstructions of all neurons within the reconstructed cartridge. From left to right: the cartridge neuron bundle, seven short visual fibres (svf), two long visual fibres (lvf) and five lamina monopolar cells (L1-L5). Shaded areas indicate the lamina layers A, B and C. These layers were based on the dendritic morphology as well as the terminals of the svfs. **D** Number and the extent of the LMC’s lateral processes (measured as the straight-line distance from the axon centre to the end point of each process). The shaded areas show the average cartridge diameter (=10 µm). **E** Synaptic connections between the cells identified based on vesicle clusters visible in the SBF-SEM data, separated into post-(top image) and presynaptic (bottom image) partners. **F** Pre - and post-synaptic connections of all LMC types of the focal cartridge, separated into synaptic partners (photoreceptors and other lamina neurons). The total number of synapses for the individual pairings is indicated in colour.

#### Image Analysis

##### 3D segmentation

We registered the image stack using TrakEM2 (Cardona et al., 2012) in Fiji (Schindelin et al., 2012). We then obtained a detailed volumetric reconstruction of the fine morphology of the different types of lamina monopolar cells, as well as the photoreceptors manually using 3Dmod (3dmod Version 4.9.12, University of Colorado). For the 3D segmentation, we selected all neurons in one central cartridge in the image stack.

To confirm that the detailed morphology of neurons in the focal cartridge extended to other cartridges, we used Catmaid (Version 2021.12.21, CATMAID Developers) to assess the number and types of lamina monopolar cells and photoreceptors, and their locations within multiple neighbouring cartridges. Further, we created skeleton reconstructions of 5 pairs of L1 and L2 LMCs to quantify the variation in the lengths and extension of their lateral processes across cartridges.

##### Synaptic connectivity

We identified all synaptic connections of the LMCs in the focal cartridge manually in 3dmod. A synaptic connection was classified as an aggregation of vesicles next to the neuron’s membrane in one cell, and there were one or more cells attached to that membrane close to the vesicle cluster (see Fig. 1E). Within these synapses we distinguished pre- and postsynaptic connections, as well as synaptic connection between the LMC and a photoreceptor or with another cell type. On L4, where we found the synapse distribution along the processes and the spread of the processes to be highly consistent, we localized the synapses on a subset of 3 processes and interpolated the total number based on these.

#### Physiology

##### Electrophysiological recordings

To perform intracellular electrophysiological recordings, the moths were fixated: their wings were cut off and their thorax and abdomen were restrained by wrapping the animals in tape such that only the head remained free. To reduce movement further, any dental wax was applied on the thorax to minimize thorax movements, and the head-thorax area to fixate the head. In addition, the proboscis was extended and fixated in front of the head, and the antennae were fixated frontally on the head. Subsequently, all scales were removed from the head capsule, and a hole was cut, extending from the dorsal rim area of the left eye to the base of the left antenna.

We used borosilicate glass electrodes (Sutter Instrument Borosilicate glass with filament, BF150-75-10) to impale the lamina neurons. The electrodes were pulled either using a P-97 micropipette puller with a 2.5 mm trough filament (Sutter instruments) and a P-1000 micropipette puller with 2.5 mm box filament (Sutter instruments). When recording, the electrode tips were filled with neurobiotin (Vector Laboratories, Burlingame, USA, 4% dissolved in 1M KCl) and backfilled with 1M KCl for a subset of the experiments and filled the tip with Alexa Fluor 568 Hydrazide (ThermoFisher Scientific) and backfilled with 200mM KCl for another set of recordings. The resistances of the pulled electrodes ranged from 80 MOhm to 120 MOhm for neurobiotin injections but were considerably higher for A568 recordings, with resistances up to 400 MOhm. After recording the neuronal responses, the neurons were dye injected by applying a constant iontophoretic current. For neurobiotin, a depolarizing current of 1.3 nA was applied for 2-3 minutes, while for Alexa568, a hyperpolarizing current of 2.5 nA was applied for 3-5 minutes.

The electrodes were controlled by a manual micromanipulator attached to a piezo control box. The recorded neural responses were pre-amplified by a headstage and then amplified by a bridge amplifier (BA-03X, NPI electronic GmbH, Tamm, Germany). Signals were monitored with an oscilloscope and an audio monitor and then digitized by using a data acquisition card (National Instruments, USB 6221). The stimuli and data acquisition were generated and controlled by a custom written Matlab script (Matlab2015a, Mathworks) used with psychtoolbox.

##### Histology

The brains containing dye injected neurons were dissected out of the head capsule and transferred to a petri dish containing 0.1M phosphate buffer (PB, pH 7.2), in which the trachea and any other loose tissue were removed from the brain. The retina of the left optic lobe was cut off with fine scissors to fully expose the lamina. neurobiotin-injected brains were fixated at 4 °C overnight in neurobiotin-fixative (4 % paraformaldehyde, 0.25 % glutaraldehyde, 0.2 % saturated picric acid in 0.1 M PB). After fixation, brains were washed 5 times for 15 minutes in 0.1 M PBS (PB containing 0.145 M NaCl) and then incubated with Streptavidin-conjugated Alexa Fluor 568 (ThermoFisher Scienfitic) 1:1000 in PBT (0.1 M PBS containing 0.3 % Triton X-100) for 3 days in darkness at 4 °C. After the incubation, the brains were washed 2 x for 30 minutes in PBT and 3 x 30 minutes in PBS. Then, brains were dehydrated in an ascending ethanol series (30 %, 50 %, 70 %, 90 %, 95 %, 100 %, 100 %) for 15 minutes each. Afterwards, they were transferred to a 1:1 mix of 100 % ethanol and methyl salicylate for 20 minutes and then cleared in pure methyl salicylate for 1h or overnight. After clearing, the brains were embedded between two coverslips in permount (Fisher Scientific, Schwerte, Germany).

The A568-injected brains were fixated in 4 % paraformaldehyde-fixative for 30 minutes at room temperature. After fixation, the brains were dehydrated, cleared and embedded as described for the neurobiotin injections.

##### Visual stimulation and data analysis of contrast and luminance response properties

During recordings, the animals were placed in front of a screen at a distance of 20 cm, such that their left eye centred on the middle of the screen. Visual stimuli were generated by a custom written Matlab script (Matlab2015a, Mathworks) used with the psychtoolbox presented at a frame rate of 120 Hz. The screen was linearly calibrated to ensure brightness consistency across the whole visual field. If applicable to the stimulus, they were corrected for the animals’ viewing perspective to that all pixels of the screen subtended the same visual angles in the field of view of the animal.

To examine the contrast and luminance response properties of the cells, we presented either a brightness increment (ON step) or decrement (OFF step), changing the screen from mid grey to white or to black, which results in a Weber contrast of 1 or −1, respectively. We presented stimuli with two durations: a short impulse of 8 ms and an impulse of 500 ms duration. The stimulus was preceded and followed by a 1 second adaptation phase presenting a mid-grey screen. Each stimulus was repeated at least 10 times during a recording.

Custom-written code in Matlab (Matlab2022b, Mathworks) was used for all following analyses. To compare the response characteristics of individual cells, we normalized the responses by the maximum amplitude and averaged all stimulus repetitions. The 8 ms impulse responses were fitted with a double lognormal function, including a positive and a negative deflection (Fig 2C, E). The fitting function included 6 parameters, which were optimized using the inbuilt Matlab function ‘fminunc’:

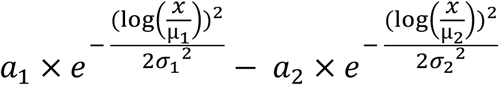

**Figure 2:**
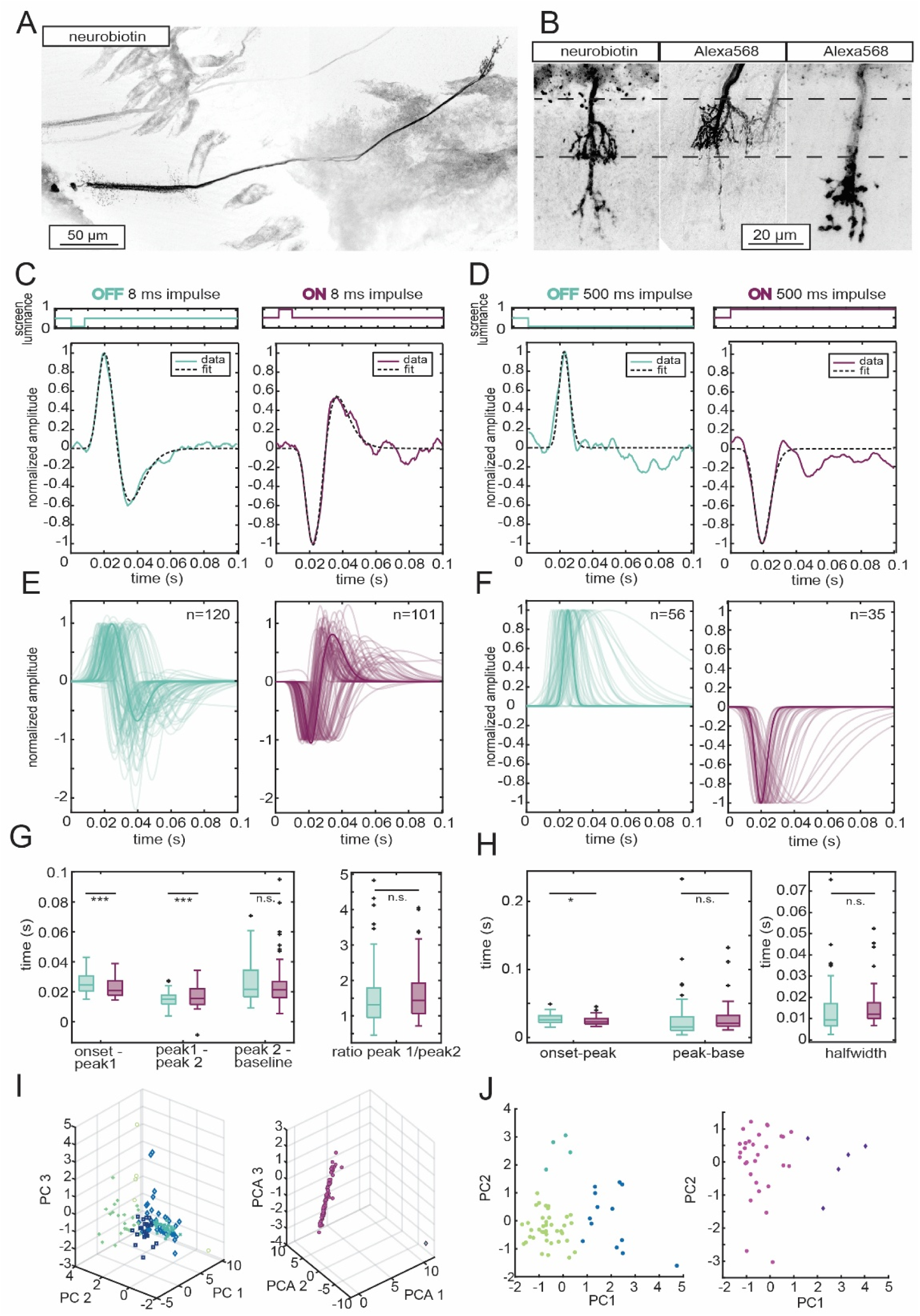
Contrast coding properties of L1 and L2. **A** Confocal scan with a 10x magnification shows a typical example of a neurobiotin injected recording: 2 neurons were stained, 2 cell bodies and a double terminal are visible. **B** Confocal scan with a 25x magnification, 3 terminals of L1 and L2 neurons. Dashed lines indicate the distal end of the medulla (top line) and the border between first and second medulla layer (bottom line). Left image: double terminal as found with neurobiotin injections with one terminal in the first medulla layer and the second terminal at the proximal side of the second layer. Middle and right image: single neuron staining with Alexa568 dye revealed single terminals in either the first or the second medulla layer. **C, D** Example recordings for OFF (left) and ON (right) impulse stimuli of **C** 8 ms and **D** 500 ms, respectively. Coloured lines (OFF = green, ON = purple) represent the raw traces, dashed line the fits. **E, F** Fits to all individual cell responses for the **E** 8 ms and **F** 500 ms ON and OFF stimulation, respectively. N-numbers represent cells, OFF and ON data can have been recorded in the same cell or in different cells. **G, H** Quantification of temporal dynamics of the fitted **G** 8 ms and **H** 500 ms ON and OFF responses. Colour code corresponds to OFF (green) and ON (purple). **G** Left: time from stimulus onset to first peak; time from the first peak to the second, inversed peak, and time from the second peak to baseline. Right: Ratio between the amplitude of the first peak and the second peak for ON and OFF. **H** Time to peak extracted from fit data: Left: stimulus onset to peak, and time from peak to baseline. Right: half-width of the fits for ON and OFF responses. All statistical analysis accounts for paired and unpaired data points (* p<0.05, ** p<0.01, *** p<0.001, n.s. not significant). **I, J** k-means clustering of the first three principal components for cell responses to **I** 8 ms and the first 2 principal components for cell responses to **J** 500 ms stimuli, respectively, based on the fitting parameters of the cell responses. Colour code remains the same (OFF= green; ON=purple colours).

To investigate potential response clusters, which could result from different cell types with idiosyncratic responses, we first reduced the parameter dimensions by performing a principal component analysis (PCA). As an input to the PCA, all parameters were transformed into z-scores to eliminate differences in units, with a mean 0 and standard deviation of 1. The results provided a measure for the proportion of variance in the data explained by each principal component (Fig S1 A, E) as well as the contribution of each parameter to each principal component (Fig S1 B, F). To reduce dimensionality, we incorporated the principal components that explained at least 80 % of the variability in the data. For both, ON and OFF impulse responses, this required 3 principal components. With these 3 principal components we performed k-means clustering to assess differences in the response properties between the recorded cells (Fig 2I). To estimate the optimal number of clusters, we calculated the silhouette score (Fig. S1 C, G), which was used as a criterion for the probability that the data was correctly assigned to the right number clusters. In general, if the average silhouette score across all points was 0.7 or higher, we considered the clustering to be reasonable. To further assess the separation between the clusters, the silhouette value for each data point within the clusters were calculated (Fig S1 D, H): numbers close to 1 indicate a clear separation between clusters, whereas low or rather negative values indicate that the data is not certainly assigned correctly to its cluster.

We further investigated differences in cell responses towards OFF and ON stimulation, by quantifying differences in timing dynamics within but also across different cells. Therefore, we compared cell response times from stimulus onset to first peak, from the first peak to the second cell deflection, and the time from the second deflection back to the baseline (Fig 2 G, left). Additionally, the ratio of the amplitudes of the first peak and the second deflection was calculated (Fig2 G, right). Statistical differences between timing dynamics for OFF- and ON cell responses were calculated by a linear mixed-effects model (lme).

The same analysis was applied to the 500 ms impulse responses with some adjustments. The data was fitted by a single lognormal function, thus fitting only the cells’ response to the onset of the stimulus:

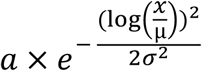

This fit resulted in 3 fitting parameters, based on which the PCA was performed. For both, ON and OFF data the first two principal components explained more than 90 % of the variance of the data (Fig S1 I, M). Thus, the subsequent k-means cluster analysis was performed on the first two principal components (Fig 2J) and statistically analysed by the silhouette method (Fig S1 K, L, O, P), as described above.

Differences in timing dynamics between ON and OFF cell responses within and also across cells were quantified by comparing response times from stimulus onset to the first peak and from the peak back to baseline. Additionally, the halfwidths of the cell responses were compared. And. as described above, statistically tested by using a linear mixed-effects model.

##### Visual stimulation and data analysis of spatial characteristics

We tested the spatial response properties of the cells by presenting a moving black bar on a white background. The bar moved from left to right and vice versa, as well as up to down and vice versa. Each bar orientation was presented 8 times, and the receptive field of the strongest responses (either horizontal or vertical) was extracted. We normalized each trial by the maximum response amplitude and averaged all normalized responses. Subsequently, we fitted the averaged receptive fields using the inbuilt Matlab function ‘isqcurvefit’ combined with a double Gaussian function, one of which had a positive and the other one a negative amplitude, creating a difference-of-Gaussian shape:

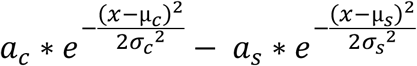

This resulted in 6 parameters per receptive field. We reduced these parameters to 5 by calculating the relation between the two amplitudes (a = a_C_/a_s_). We further reduced the dimensionality of the data using a PCA (as described above). The first 4 principal components explained more than 80% of the data (Fig S2 A). Based on these we then evaluated the optimal number of clusters for k-means clustering, using the silhouette score and the gap statistic to evaluate the separation of clusters (Fig S2 C, D).

##### Spatial modelling

We used the distribution of L1 and L2 post-synapses to predict the spatial response properties of these neurons. For all comparisons to physiological responses, we relied on data from Stöckl et al. (2020), as this dataset included both photoreceptor and LMC recordings obtained with the same experimental setup. Photoreceptor responses are required for response predictions (see also the modelling in Stöckl et al. 2020), and therefore using data acquired under identical conditions is critical to minimize variance introduced by stimulus geometry and recording position. The L1 and L2 post-synapses were separated into the three lamina layers A, B, and C (Fig. 4A), and histograms of their distances to the cartridge centre were calculated (Fig. 4B, C for L1 and L2, respectively). For subsequent LMC filter predictions, we used the average of L1 and L2 layers A, B, and C histograms.

Because we compared anatomical predictions of spatial processing with physiological measurements of LMC responses, we required a conversion from the distance between neighbouring processing units (cartridges) to visual angles, which are the units in which physiological spatial response properties are quantified. To obtain this conversion factor, we fitted a Gaussian distribution to the average layer B synapse histogram and minimized the squared distance of this distribution to photoreceptor responses in bright light (100 cd/m²). Importantly, this distribution was fitted to photoreceptor, rather than LMC, responses, because the short processes integrate photoreceptor input, whereas LMC responses reflect both photoreceptor input and lateral inhibition (likely mediated by longer lateral processes) under bright light conditions. The resulting conversion factor was 1.642° per cartridge-to-cartridge distance, closely matching previous optical measurements in the lateral eye by Warrant (1999), taken from the same region of the lamina tissue used for our anatomical investigation.

We then constructed LMC spatial processing filters based on the synapse distributions of the short layer B processes, combined separately with the longer processes of either layer A or layer C. This allowed us to assess whether layer A or C provided better fitted for the following spatial processing tasks: lateral inhibition and spatial summation. To construct lateral inhibition filters, the synapse distribution of layer A or C was subtracted from that of layer B, whereas for spatial summation the two were added. We did not introduce additional weighting factors but scaled solely by the number of synapses in each respective layer (i.e. the amplitude of each synapse distribution). These spatial filters were then convolved with the physiologically measured photoreceptor responses (see also Stöckl et al. 2020): inhibition filters with photoreceptors recorded in bright conditions, and summation filters with photoreceptors recorded in dim conditions (Fig. 4D).

The goodness-of-fit of the spatial response predictions, obtained by convolving the spatial filters with the photoreceptor responses, was then compared to physiologically measured LMC responses in both dim and bright light using Euclidean distances (see also Stöckl et al. 2020) (Fig. 4E, F). We statistically compared Euclidean distances between predictions and LMC responses with distances across LMC responses themselves (as a measure of variation), using Kruskal–Wallis tests in bright (χ²: 8.22, df:2, p_filterA-filterC_: 0.021, p_filterA-LMC_: 0.028, p_filterC-LMC_: 0.578) and dim (χ²: 15.37, df:2, p_filterA-filterC_: 0.85, p_filterA-LMC_: 0.003, p_filterC-LMC_: 0.032) conditions, respectively (Fig. 4E,F, top of each panel). We further compared the goodness-of-fit between the layer A and layer C filters using paired signed-rank tests in bright (zval: 3.29, signedrank: 146, p_filterA-filterC_: 0.008) and dim (zval: −2.656, signedrank: 29, p_filterA-filterC_: 0.0079) conditions (Fig. 4E, F, bottom of each panel).

## Results

### A cartridge in the hawkmoth lamina contained previously unknown cells

We used serial block-face scanning electron microscopy (SBF-SEM) (Denk & Horstmann, 2004) to reconstruct the fine structure of the neurons in one lamina cartridge of the hummingbird hawkmoth *Macroglossum stellatarum*.

The SBF-SEM data revealed a retinotopic organization of lamina monopolar cells (LMCs) and photoreceptor axons, which formed tight cartridges that could readily be separated throughout the full extent of the imaged tissue (Fig 1B). Each cartridge contained nine photoreceptor axons (Fig 1C). Two of which were long visual fibres, extending into the medulla, whereas the remaining seven photoreceptors were short visual fibres that terminated 20 µm before the proximal end of the lamina. The 3D reconstruction of the photoreceptors revealed that they did not have any lateral processes. In addition to the photoreceptor axons, each cartridge contained five morphologically distinct lamina monopolar cells (Fig 1C). This finding was surprising, as previous hawkmoth LMC classifications based on Golgi stainings described only four different types (Stöckl et al., 2016b). This discrepancy might have arisen since two of the cell types identified in our reconstructions closely matched the morphology of a single previously described type (L1 and L2, matching type II). In addition, we discovered one cell type that had not been reported before (L5).

The five LMCs showed extensive lateral branching (Fig 1D), which delineated three lamina layers: the distal layer A, middle layer B and proximal layer C. The axon terminals of the short visual fibres demarked the border between layers B and C, and the branching patterns of the LMCs defined the border between layers A and B. Of the five LMCs, the two cells with the largest axon diameters (∼ 4 µm) had similar lateral branching patterns: they had long lateral processes in layers A and C, which left the home cartridge and extended up to three to four neighbouring cartridges. In the central layer B, they had mostly short lateral processes which remained in the home cartridge (Fig 1 C, D). Both cells had close contact to each other via their axon membranes throughout the full extent of the lamina (Fig. 1B, cell IDs: L1 and L2). The LMC with the third biggest axon diameter (∼ 2 µm) had few lateral processes in layers A and C, and a regular spacing of short lateral processes in layer B, which were restricted to the home cartridge (Fig. 1C, cell ID: L3). While the axons of these three cell types remained in close contact with each other throughout the whole lamina, the two remaining cells were located further outside the cartridge boundaries (Fig 1 B). Their axons were considerably smaller in diameter (< 2 µm) throughout the whole lamina. One of the cells (cell ID: L4) had regular, mid length, branched lateral processes in layers A and B, but no processes in layer C. The fifth cell (cell ID: L5) had a few lateral processes in layers A and C, which remained in the home cartridge, and were bulky and filled with vesicles (Fig S3 A, B). This cell type did not have processes in layer B. We found the same number and arrangement of cell types in all surrounding cartridges (35 in total) across the whole image stack.

### A novel LMC classification for hawkmoths

To identify LMC types, we relied on the neuron’s morphology and their synaptic connectivity, and compared these features with the existing literature on the two insect species with 3D morphology and connectome reconstructions: the fruit fly *Drosophila melanogaster*, and the swallowtail butterfly *Papilio xuthus*. To this aim, we identified pre - and post-synaptic connections of all five LMCs in the focal cartridge to both photoreceptors and other neurons (Fig 1E). As in other insects, we termed the pair of neurons with the largest axon diameters and a central placement in the cartridge L1 and L2 (Boschek, 1971; Ribi, 1978, 1987; Shimohigashi & Tominaga, 1999). Their connectivity matched L1 and L2 cells across other insect species, as they had the highest number of synapses with photoreceptors out of all LMC types (Fig 1F; Fig S3C, D) (Rivera-Alba et al., 2011). While L1 and L2 are morphologically indistinguishable in the lamina of flies (Fischbach & Dittrich, 1989), their connectivity differs: L1 has more presynaptic connections compared to L2 in fruit flies (Rivera-Alba et al., 2011), as well as in *Papilio xuthus* (Matsushita et al., 2022). We used this indicator to identify L1 and L2 in our 3D reconstructions (Fig. 1F).

Based on the brush-like structure of its lateral processes, its axon diameter and the location within the cartridge, we classified the third LMC type as L3. This cell received its main input from photoreceptors of the home cartridge (Fig S3D), which aligns with findings in other insect species (Matsushita et al., 2022; Ribi, 1975a; Ribi, 1975b, 1987).

The fourth identified LMC, by means of exclusion of the other cell types termed L4, differs distinctly from L4 LMCs in *Drosophila* (Fischbach & Dittrich, 1989), bees (Ribi, 1975a), ants (Ribi, 1975b) and *Papilio aegus* butterflies (Ribi, 1987), which possess long lateral processes in layer C only. It was, however, previously observed in several studies of hawkmoths (Stöckl et al., 2016b; Strausfeld & Blest, 1970) and does bear some resemblance to L4 in *Papilio xuthus* (Matsushita et al., 2022). While morphologically very different, its connectivity, with nearly no postsynaptic connections to photoreceptors (Fig S3 E), does align with L4 in the two species with known connectivity, *D. melanogaster* (Rivera-Alba et al., 2011) and *P. xuthus* (Matsushita et al., 2022).

The LMC with processes in layers A and C only, is morphologically equivalent to *Drosophila* L5 (Fischbach & Dittrich, 1989) and in desert ants (Ribi, 1975b). In both species, L5 possess only few lateral processes in layer A and C. However, in *Drosophila*, its connectivity is only postsynaptic (Rivera-Alba et al., 2011), whereas we found mainly presynaptic connections (Fig S3A, B).

### Functional differences in L1 and L2 contrast and luminance coding

Having established a new classification of lamina cell types, and identified L1 and L2 neurons, we next investigated whether functional specializations existed between these two cell types, which have previously been described in fruit flies (Joesch et al., 2010). We therefore investigated imbalances in their responses to light increments and decrements, as well as differences in their responses to contrast stimuli, as *Drosophila* L1, which initializes the ON pathway, responds with more sustained responses to contrast steps than L2, which initializes the OFF pathway (Clark et al., 2011; Joesch et al., 2010). To quantify the response characteristics of hawkmoth L1 and L2, we recorded intracellularly and injected neurobiotin to characterize the recorded neurons (Fig 2A).

The dye injections after recordings confirmed that we were recording from L1 and L2. Interestingly, in all neurobiotin injections, we found at least 2 cells in the same cartridge to be strongly labelled, suggesting that L1 and L2 were coupled via gap junctions, which neurobiotin is known to pass (Fig 2A). In these double-cell labelled brains, we found terminals in two distinct layers of the medulla, the first one at the distal end of layer 1 and the second one at the distal end of layer 2 (Fig 2A, B left image). This aligns the features of L1 and L2 in fruit flies, which are coupled via gap junctions (Joesch et al., 2010), and terminate in layers 1 and 5 (L1) and 2 (L2). To separate the individual LMCs, we used injections of Alexa568 dye, which has a larger molecular structure than neurobiotin and therefore passes gap junctions less frequently. Thereby, we could isolate the two terminals in layers 1 and 2 (Fig 2B mid and right image), confirming that only single cells were labelled by registering only single cell bodies, which demonstrates that L1 and L2 terminate in distinct layers in the medulla, as they do in other species.

While the higher molecular size of the Alexa 568 dye allowed for a morphological distinction of L1 and L2, it also caused the electrodes to clog more easily during the recordings and injections, which made this method distinctly less reliable than neurobiotin injections. Therefore, we could not use this method to screen for functional response differences between the two cell types. We therefore investigated whether the recorded response of all cells identified as L1/L2 formed two distinct clusters that would delineate the two types.

We began our functional investigation with characterising the contrast and luminance coding of light increments and decrements. L1/L2 responded with a depolarization to OFF light steps (Fig 2C left) and a hyperpolarization to ON steps (Fig 2C right). To brief, 8 ms impulses, the cells responded with a quick response peak, followed by a membrane potential reversal after the offset of the stimulus (which represented an inverse contrast step to the neurons). To investigate potential differences in response amplitudes to ON and OFF steps between cell types, as well as differences in the temporal response characteristics, the responses to ON and OFF steps were fitted with a lognormal function (Fig 2E). Based on these fits, we extracted temporal response characteristics, such as the time from stimulus onset to the peak response, as well as time to membrane potential reversal after stimulus offset, the time to return to baseline, and the ratio between amplitude at the onset and offset of the stimulus (Fig 2G). The quantification of these temporal dynamics revealed that the variability across cells (SD = 0.0059 s) was larger than the variability within cells (SD = 0.0025 s) (Fig. 2E, F). Therefore, most variability in cell responses can be explained by differences between different cells rather than within cells. Further, we found that the first peak was on average 3 ms faster for the ON stimulus (p= 10^-15), whereas the second peak occurred on average 2 ms faster during the OFF stimulus (p= 0.0002) (Fig. 2G). This suggests that the cells hyperpolarized faster than they depolarized. The timing dynamics from the second peak to baseline, as well as the ratio between first and second amplitude did not reveal any significant differences between OFF- and ON-stimulation (Fig. 2G). Additionally, a cluster analysis was applied to the temporal parameters after dimensionality reduction. Reviewing the data in the principal component space (Fig 2I) showed that it was distributed continuously rather than delineating into two clusters, as would be expected if L1 and L2 had different response properties. While a statistical evaluation suggested the optimal number of clusters was five for OFF responses (Fig S1 C), the silhouette score remained lower than our set criterion of at least 0.7 (see Methods). Additionally, many data points were assigned to the clusters with low certainty (Fig S1 D). Reviewing the parameters of the ON responses in principal component space showed a continuous distribution with one outlier (Fig 2I, right). The optimal number of clusters was two supported by a high average silhouette score (Fig S1 G), however reconciling the silhouette values for the individual datapoints showed that the outlier was the only one data point assigned to the second cluster (Fig 2I right, Fig S1 H). Thus, we did not find that the neurons’ responses to ON and OFF steps clustered into two distinct groups that would represent L1 and L2.

We next tested 500 ms ON and OFF steps (Fig 2 D), as fruit fly L1 and L2 differ in their temporal response characteristics to such contrast stimuli, with L1 responding with a combination of transient and sustained responses, thus encoding both contrast and luminance, and L2 encoding primarily the contrast information in highly transient responses (Clark et al., 2011). To these long contrast steps, most neurons responded with a rapid decrease or increase in membrane potential (Fig 2 F), depending on the contrast polarity, which went back to baseline about 15 ms (OFF) and 20 ms (ON) later (Fig 2 H). While most neurons showed such a quick return to baseline, some took considerably longer, resulting in more transient cell responses. Quantifying differences in the timing dynamics towards ON and OFF stimulation revealed that the time from stimulus onset to first peak was on average 2 ms faster in the ON condition compared to OFF (p= 0.013), while there were no significant differences in the time from peak to baseline or in the halfwidth of the responses (Fig 2H). This supports the notion that hyperpolarization was faster than depolarization in these cells. To assess whether these responses fell into two distinct categories, we performed a cluster analysis similar to the 8 ms contrast steps. As for those, assessing the response parameters in principal component space showed a continuous distribution for both ON and OFF steps (Fig 2J). The optimal number of clusters for OFF and ON was three and two (Fig S1 K, L & O, P), with only very few cells represented in the second or third clusters, which did not fit the expectation of two equally distributed response groups corresponding to L1 and L2, respectively.

Across short and long contrast steps with brightness increments and decrements, we did not observe two distinct response clusters which we would have expected if L1 and L2 had clearly distinguishable contrast response patterns.

### Spatial properties of L1 and L2

We next investigated whether there were differences in the spatial morphology of L1 and L2, and the corresponding spatial response properties. This question was initiated by the morphological differences of the L1 and L2 neuron that were fully 3D reconstructed in the focal cartridge (Fig. 1C, D): the focal L2 neuron had distinctly longer lateral processes than L1. A skeleton reconstruction of the lateral processes of L1 and L2 in four additional cartridges revealed, however, that these differences in lateral extent were not consistently aligned with neuron type, but varied from cartridge to cartridge (Fig. 3A, B). This suggests that process length is not consistently different between L1 and L2, but might be rather randomly varying, possibly as a consequence of their development.

**Figure 3.**
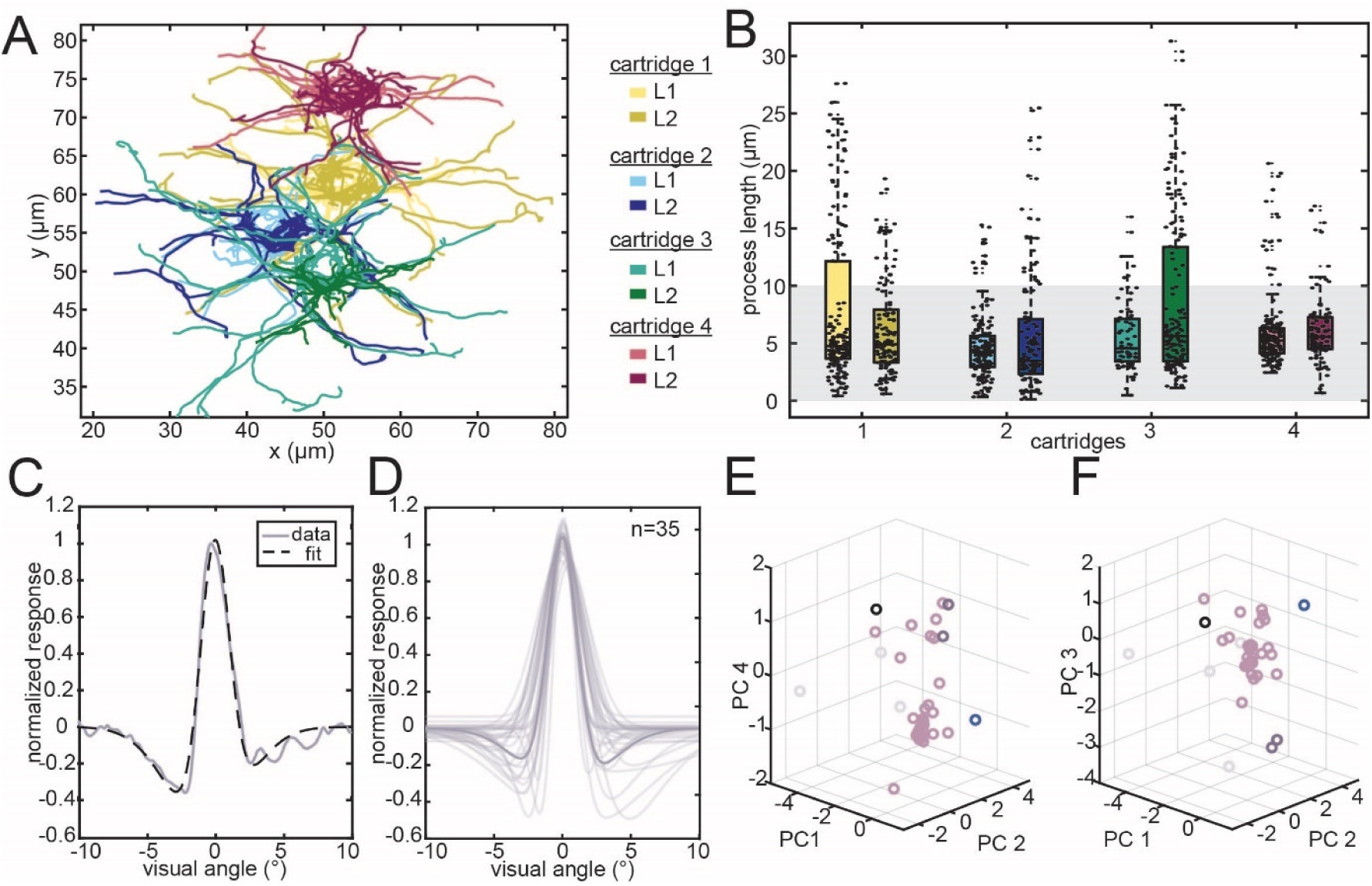
Spatial processing properties of L1 and L2. **A** Skeleton reconstruction of L1 and L2 neurons in 4 different cartridges as a top view, showing the lateral extents of the processes in lamina layer A. **B** Comparison of the process length of L1 and L2 processes in layer A across the 4 cartridges. **C** Example recording of an L1 / L2 receptive field. Coloured line shows the raw trace, dashed line shows the fit. **D** Difference-of-Gaussian fits of all individual receptive field measurements recorded from 35 different cells. **E** k-means clustering of the first 2 principal components and principal component 4, different colours represent the 5 clusters **F** k-means clustering as seen in E but showing the first 3 principal components.

Correspondingly, we analysed the spatial response properties of the neurons with intracellular recordings. In bright light intensities, the recordings revealed receptive fields with a centre-surround structure (Fig 3C). We therefore fitted the receptive fields with a difference of Gaussians (Fig 3D), which revealed an average centre half-width of 2.6 degrees, and an average half width of the inhibitory surround of 7.7 degrees. Clustering of the receptive field parameters similarly to the temporal properties revealed that the optimal number of clusters was either one or five (Fig S2 C), reviewing the data in the principal components space for the first 4 principal components does not show a clear separation of the data (Fig 3E, F) again suggesting no distinct differences in spatial responses between L1 and L2.

### Synaptic distribution reveals spatial processing strategies

Since L1 and L2 in hawkmoths play a key role in spatial processing, both lateral summation in low light (Stöckl et al., 2020) and as demonstrated here, lateral inhibition in bright light (Fig. 3C), we next investigated whether the distribution of pre - and post-synapses across layers A, B and C would reveal how these processing strategies delineate across the morphology of the neurons (Fig 4A). In layer B, the short lateral processes contained within the home cartridge carried only post-synapses to photoreceptors (Fig 1F), while the long processes in layer A and C formed synaptic connections with other cell types (Fig 1F). This synaptic architecture provides important insights into the mechanisms of spatial processing in L1 and L2: the centre of the receptive fields of the neurons matched the spread of the post-synapses in layer B (Fig 4B, C), suggesting that these were formed by the information which the short processes of the neurons integrated, while lateral processing is confined to layers A and C. Importantly, in these layers, the post-synapses were present on processes that extended outside of the home cartridge, whereas pre-synapses were found only within a range of 10 µm from the axon on the long processes in layers A and C (Fig. S4). If these mitigate spatial summation or lateral inhibition, the extent of this processing would therefore only be determined by the length of the lateral processes of other L1 and L2 neurons, as there would be no synaptic transmissions between the long processes of L1 and L2 outside of the respective home cartridges of both pre- and post-synaptic partners.

**Figure 4.**
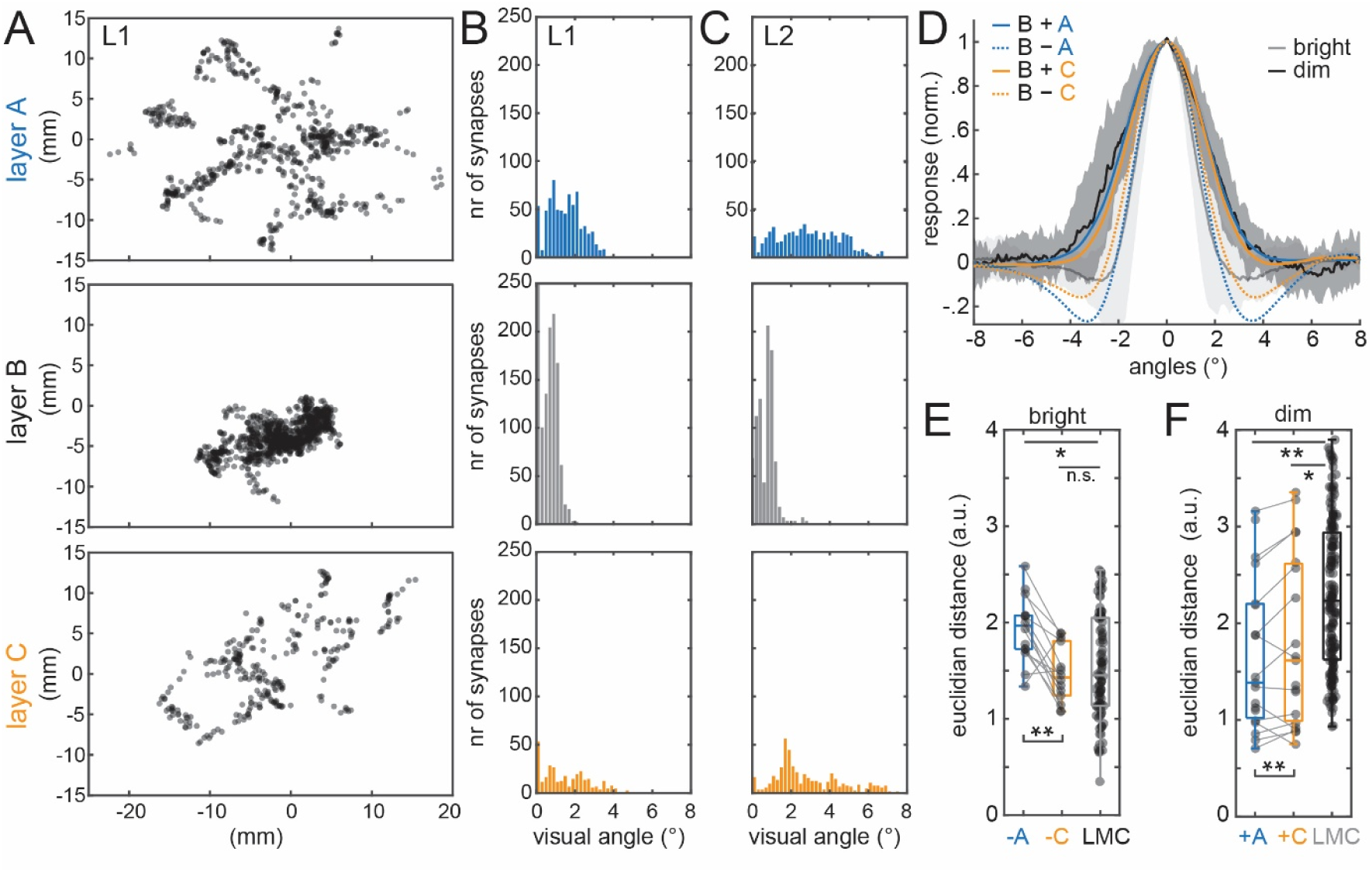
Spatial processing model based on synaptic distributions of L1 and L2. **A** Post-synapses of L1, in layers A, B and C (see Fig. 1C for layer definition, see Fig. S4B for L2). **B, C** Post-synapse histograms for the fully reconstructed L1 and L2 neurons. **D** Lateral inhibition and spatial summation response predictions, based on either subtracting or adding layer A or C to layer B synapses, respectively (see Spatial Modelling Methods for details). Goodness-of-fit between physiological lamina monopolar cells response and response predictions based on either layer A or C contributions to spatial filters (see **D**), providing lateral inhibition in **E** bright, and spatial summation in **F** dim light. Euclidian distances across the LMC responses provide a measure of the inherent data variation (LMC). Kruskal-Wallis-tests with Tukey–Kramer corrected post-hoc tests were performed to assess differences between response predictions and variation in data (results depicted on top of the panels). Paired signed-rank tests were performed to assess differences between the response predictions with layer A or C contributions (results depicted at the bottom of the panels). * p<0.05, ** p<0.01, *** p<0.001, n.s. not significant.

To test whether lateral inhibition and spatial summation are likely performed in either layer A or C, respectively, we used the distribution of L1 and L2 post-synapses (Fig 4B, C) to construct spatial filters, predicting LMC responses formed by inhibition and summation in layers A and C, respectively, in addition to the integration of photoreceptor responses by layer B synapses (for modelling details, see Methods) (Fig. 4D). These predictions revealed that the subtractive combination of layers A and B synapses had significantly lower fitting errors than those of layers C and B to physiologically measured LMC receptive fields containing lateral inhibition in bright light (Fig. 4E), while the additive combination of layer A and B synapses had a significantly lower fitting error to LMC receptive fields revealing spatial summation in dim light (Fig. 4F). These results suggest that different spatial processing functions might be delineated into separate layers of the lamina: summation into the distal layer A and lateral inhibition into the proximal layer B.

## Discussion

Using serial block-face scanning electron microscopy (SBF-SEM), we provide a novel classification of lamina monopolar cell types in a hawkmoth species, based on 3D-morphological features and synaptic connectivity. Our analysis revealed five lamina monopolar cells (LMCs) per cartridge, which we assigned to the established L1-L5 types homologous to other insect species.

### Reconciling lamina classifications across species

In order to extrapolate and compare functional characteristics of LMCs across insect species, homology of these cell types is required. Establishing homology based on the neurons’ morphology in the lamina and medulla seems challenging. First, different insect species are reported with different numbers of LMCs per cartridge: most species across insect groups possess five LMC types (Diptera: fruit flies (Fischbach & Dittrich, 1989), Hymenoptera: desert ants (Ribi, 1975b), Coleoptera: fireflies (Ohly, 1975), Lepidoptera: skipper butterfly (Shimohigashi & Tominaga, 1999), hawkmoths, and in some cartridges of the Japanese yellow swallowtail (Wakita et al., 2024). Other species, even in the same insect groups, were only reported to have four (Hymenoptera: honey bee (Ribi, 1975a), tropical sweat bee (Greiner et al., 2004b), Lepidoptera: orchard swallowtail butterfly (Ribi, 1987)). It is possible that a fifth LMC is present in all or some of the latter species, and was missed because two LMC types share a similar morphology, as was the case for the hummingbird hawkmoth we investigated here.

Second, some morphological features are consistent across species, while others can differ vastly. For example, brush-like processes on one side of the axon associated with L3 have been found in ants (Ribi, 1975b), honeybees (Ribi, 1975a), the fruit fly (Fischbach & Dittrich, 1989), firefly (Ohly, 1975), the orchard swallowtail butterfly (Ribi, 1987) and the hummingbird hawkmoth (Stöckl et al., 2016b), Fig. 1C)). However, neurons with this characteristic morphology are absent in other species, such as sweat bees (Greiner et al., 2004b), and the Japanese yellow swallowtail (Wakita et al., 2024). This morphological variation is striking, particularly since it is also present across relatively closely related insect species. Another example of diverging morphology is L4, which possess lateral processes that only extend within lamina layer C in fruit flies (Fischbach & Dittrich, 1989) and in the orchard swallowtail butterfly (Ribi, 1987), but not in the hummingbird hawkmoth, or the Japanese yellow swallowtail (Wakita et al., 2024). The axon diameter, and position within a cartridge is also highly consistent for some neuron types, particularly L1 and L2, which are typically at the center of a cartridge and have the largest axon diameters of all LMC types in house flies (Pyza & Meinertzhagen, 1995), fruit flies (Fischbach & Dittrich, 1989), the honeybee (Ribi, 1975a), the hummingbird hawkmoth (shown here), but not in the Japanese yellow swallowtail butterfly (Wakita et al., 2024).

Another commonly used approach to classify LMC types across insects is the termination layer of the LMC axon terminals in the medulla, which provided the initial naming convention for fruit fly LMCs (Fischbach & Dittrich, 1989). However, while different LMC types consistently project to distinct medulla layers within a species, the termination layers themselves are not conserved across insect groups: across the known lamina neuron types in Hymenoptera, L1 terminates in layer 1 in honeybees and desert ants, and in layer 2 in sweat bees, while this relation is swapped for L2. L3 and L4 also do not terminate in the same layers across honeybees, sweat bees, and desert ants (Ribi, 1975b), (Greiner et al., 2004b; Ribi, 1975a).

Thus, the morphology of the neurons in the lamina and medulla does not provide sufficient information to establish homology. We suggest that combining a neurons’ synaptic connectivity with their 3D morphology makes it possible to identify functional homologies, across Lepidoptera (*Papilio xuthus* (Matsushita et al., 2022), *Macroglossum stellatarum* in this study) and Diptera (*Drosophila melanogaster* (Rivera-Alba et al., 2011)). Despite their striking morphological differences, L1, L2 and L3 in these species are the main post-synaptic partners to photoreceptors. L4 has little to no postsynaptic connections to photoreceptors, but receives most of its inputs from other (lamina) cells. L5 is mainly pre-synaptic in all 3 species.

The similarities in connectivity could be the result of an evolutionarily conserved wiring scheme of the lamina across insect species, as is also known from other brain areas (Heinze, 2024), while the species’ specific differences in LMC morphology reflect adaptations to specific processing requirements, in terms of luminance adaptation, spatial or colour processing.

### Functional homology of the main relay neurons of the motion pathway

Leveraging the proposed homology between LMCs in the hummingbird hawkmoth and fruit fly, we investigated whether their LMCs share functional characteristics. We focused on neurons with the best physiological characterization, L1 and L2, which are the main contributors to the wide-field motion vision pathway in fruit flies (Borst, 2014; Borst et al., 2020; Egelhaaf, 2023).

They separate visual information in light increments (ON pathway), and decrements (OFF pathway) at their terminals in the medulla (Behnia et al., 2014; Joesch et al., 2010). Even though L1 and L2 are known to be interconnected via gap junctions in fruit flies (Joesch et al., 2010), L2 in fruit flies shows distinct selectivity to OFF signals in its terminal calcium responses, suggesting that signal rectification happens in the cell. Similar to the fruit fly, we demonstrated that in hummingbird hawkmoth L1 and L2 are likely linked via gap junctions and their terminals also end in layers 1 and 2 in the medulla (Fig 2B).

We attempted to identify which of the terminals was associated with L1 and L2, respectively, as identified based on the SBF-SEM data. However, since the 3D morphology of L1 and L2 was highly similar, and the extend of the lateral processes could also not be used as an indicator of neuron identity, as it varied more strongly within than across L1 and L2 cell types, assigning identity of the recorded and subsequently labelled neurons via morphological means was impossible. In our recordings of hawkmoths L1/L2, we also did not observe two distinct response clusters to ON and OFF visual stimuli, which would represent distinct response characteristics to light increments and decrements, suggesting that both morphologically, and in terms of their response polarity to contrast steps, L1 and L2 were not distinguishable.

We therefore planned to establish neuron identity via their temporal response characteristics, akin to fruit flies (Clark et al., 2011). However, in the hummingbird hawkmoth, we did not find two response clusters that would indicate that L1 and L2 had different temporal response properties, encoding different proportions of contrast and luminance. Both responded highly transiently, as reported previously (Stöckl et al., 2020), indicating that they primarily encode contrast information, but not luminance (Fig 2 F). Thus, if hawkmoths L1 and L2 initialized ON and OFF pathways in the motion circuitry similar to fruit flies, both pathways would be provided with highly similar response properties in terms of increment and decrement amplitude and contrast coding by the two cell types.

### Reconciling spatial and contrast processing in motion processing

Our identification of L1 and L2 reconciled two processing features in insect motion processing: segregation of light information into increments and decrements (ON and OFF pathways), and spatial integration. It has been suggested anatomically (Greiner et al., 2005; Stöckl et al., 2016b) in various insect species, and demonstrated physiologically in the hummingbird hawkmoth (Stöckl et al., 2020) that the lateral processes of LMCs that extend into neighbouring cartridges are capable of integrating visual information in dim light, to improve the signal to noise ratio, and thus contrast sensitivity, of motion responses (Stöckl et al., 2017; Stöckl et al., 2016a). How such spatial processing applies to the parallel ON and OFF arms of the motion pathway was not clear, because the neurons previously classified as L1 and L2 in hawkmoths had very different lateral extents (Stöckl et al., 2016b), suggesting that spatial processing was applied asymmetrically to the ON and OFF pathways.

With our novel LMC classification in hawkmoths we demonstrate that L1 and L2 supply the same potential for spatial processing, in terms of lateral extent, to the ON and OFF arms of the motion pathway. Our physiological investigation of the neuron’s receptive fields supports this notion, as we found no evidence for two distinct clusters in the spatial receptive fields (Fig. 3C, Fig S2). Thus, assuming that L1 and L2 initiate On and OFF pathways in hawkmoths, we demonstrate that spatial information is transmitted symmetrically into both arms. Whether this is also the case in other insects with asymmetric spatial morphologies in L1 and L2 (Greiner et al., 2004b; Ribi, 1975a; Ribi, 1987) remains to be tested.

### Synaptic architecture for spatial processing

In addition to the previously described spatial integration of information in dim light, which dynamically broadens receptive fields of L1/L2 in the hummingbird hawkmoth (Stöckl et al., 2020), in our current recordings in bright light we observed another spatial processing phenomenon, which is observed in many peripheral visual neurons in bright light: lateral inhibition (Fig 3C,D) (Hartline, 1969; Kramer & Davenport, 2015; Mimura, 1976; Zettler & Jarvilehto, 1972), thought to enhance lateral contrast for improved spatial acuity, particularly in bright light (Priebe & Ferster, 2008; Strausfeld & Campos-Ortega, 1977).

L1 / L2 processes in layer C contained considerably more pre-synapses than post-synapses compared to layer A. Together with short visual fibres of the photoreceptors terminating in layer B, this suggests that primary input processing occurs before layer C. Instead, layer C appears to be well suited for lateral inhibition.

Having reconstructed the synaptic distributions of L1/L2, we provide a mechanistic explanation for these two spatial processing functions of lamina monopolar cells, based on the density and distribution of pre- and postsynaptic connections of L1 and L2. Notably, we find that almost all postsynaptic connections to photoreceptors in L1 and L2 are concentrated in layer B, with no synaptic connections to other neurons in this layer. This suggests that layer B is the primary site for sampling photoreceptor input. Importantly, our synaptic data answers an open question concerning the extent of spatial integration: it was previously shown that the extent of the recorded receptive fields containing signatures of spatial summation in low light matched the extend of the long lateral processes of L1 / L2 in hummingbird hawkmoths. However, assuming connections between these lateral processes, the receptive fields should have been considerably wider. Our data, however, demonstrates, that there are no (chemical) pre-synapses on the lateral processes beyond the extent of the home cartridge (Fig 1F). Thus, the capacity for spatial summation (assuming no involvement of additional neurons in the lamina) in hummingbird hawkmoths is constrained by the extent of their lateral processes, which explains the close match between their spatial receptive fields in dim light, and their process distributions observed previously (Stöckl et al., 2020).

Lateral processing in hummingbird hawkmoth L1 and L2 thus concentrates in layers A and C, separated by the main input layer. Our connectivity data provides a strong hypothesis for how spatial processing functions segregate across these layers: we propose that the lateral processes in layer A perform spatial summation of information in low light, while those in layer C perform lateral inhibition. (Fig. 4).

Based on these findings, we propose that the different lamina layers support distinct connectivity and functional roles. Specifically, our findings support the existence of a layer-based mechanism for spatial processing in the hawkmoth lamina, separating summation and lateral inhibition in a similar – albeit much simpler – architecture as the vertebrate cortex (Luo, 2021). Our results not only reinforce the notion that morphology and connectivity provide strong indicators of cell function, but they also allow us to formulate new hypotheses about the homology of functional circuitry across insect species.

## Author Contributions

Conceptualization: A.S.; Methodology: A.S., K.A., R.B.; Validation: R.B.; Formal analysis: R.B.; Investigation: R.B.; Resources: A.S., K.A.; Data curation: R.B.; Writing - original draft: R.B.; Writing - review & editing: R.B., K.A., A.S.; Visualization: R.B.; Supervision: A.S.; Funding acquisition: A.S., K.A.

## Acknowledgements

We would like to thank Christian Stigloher for excellent suggestions regarding 3D tracing tools, and Nadine Kaiser and Madita Naumann for supporting the 3D neuron reconstructions. We thank Stanley Heinze and Vallentin Gillet for making their catmaid environment available for our data analysis.

## Funding

We acknowledge funding to A.S. by the German Research Council (STO 1255 2-1), the Young Scholar Fund of the University of Konstanz, and the European Research Council (101116305 DynamicVision ERC-2023-STG), and to K.A. by the KAKENHI of the Japan Society for the Promotion of Science (JSPS) (23K26914).

## Data Availability

preliminary data link: https://figshare.com/s/70a7b36da8b7dcbe0040

## Competing Interest Statement

The authors declare no competing interests.

## Supplementary Figures

**Figure S1:**
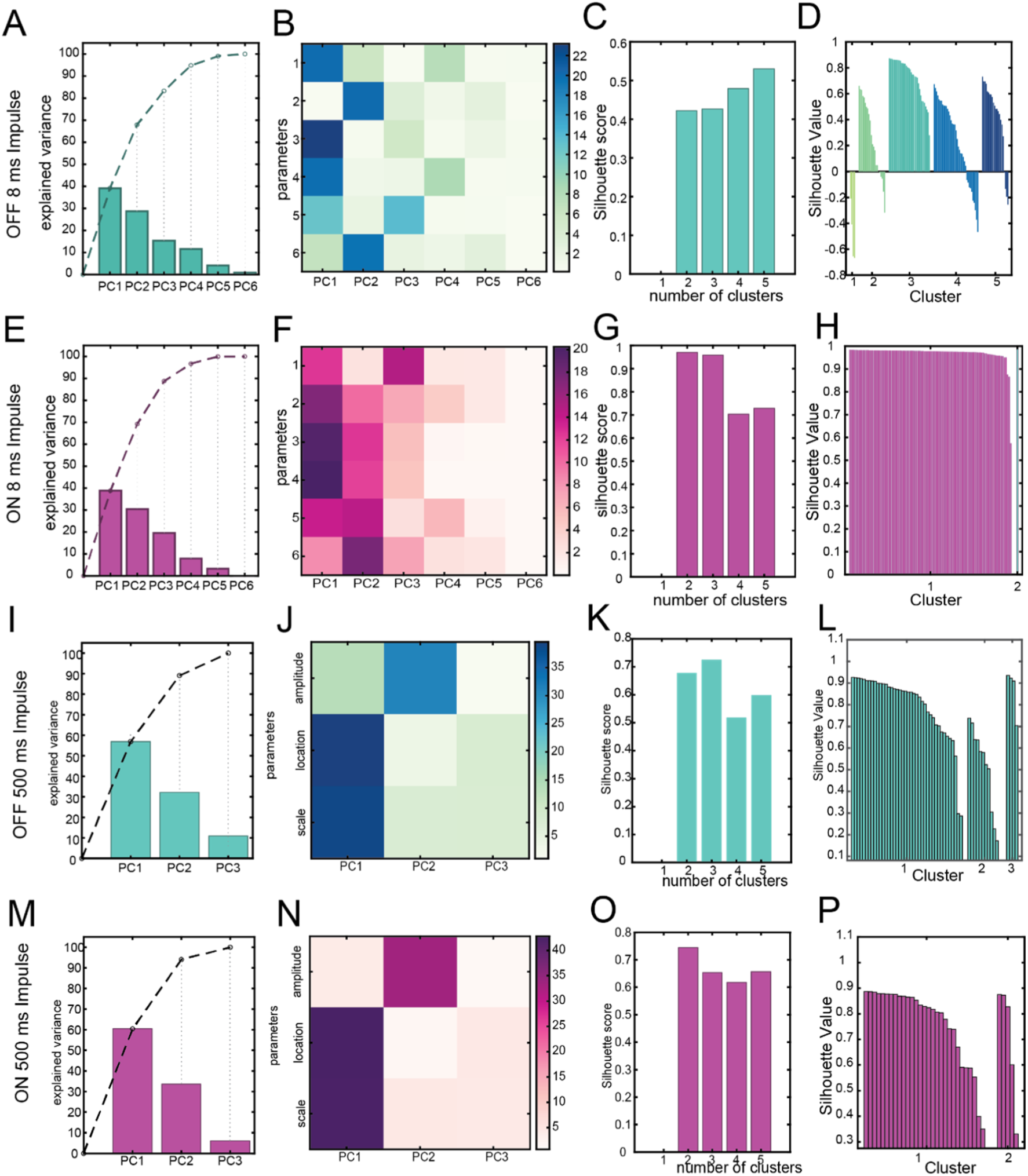
**A** Cumulative explained variance of the individual principal components which are based on the fit data of 8 ms OFF impulse responses (Fig. 2). **B** The heatmap shows which fitting parameters contribute most to which of the principal components. **C** The silhouette score gives a measure for the optimal number of clusters. Values under 0.6 are considered very low and highlight a high uncertainty. **D** The silhouette plot for the suggested number of clusters. Negative values indicate a high chance of data points being assigned to the wrong cluster. Similarly, low positive values indicate a high uncertainty in cluster assignment. **E-H** same analysis as A-D for 8 ms ON responses, **I-L** for 500 ms OFF impulse responses, and **M-P** for 500 ms ON responses.

**Figure S2:**
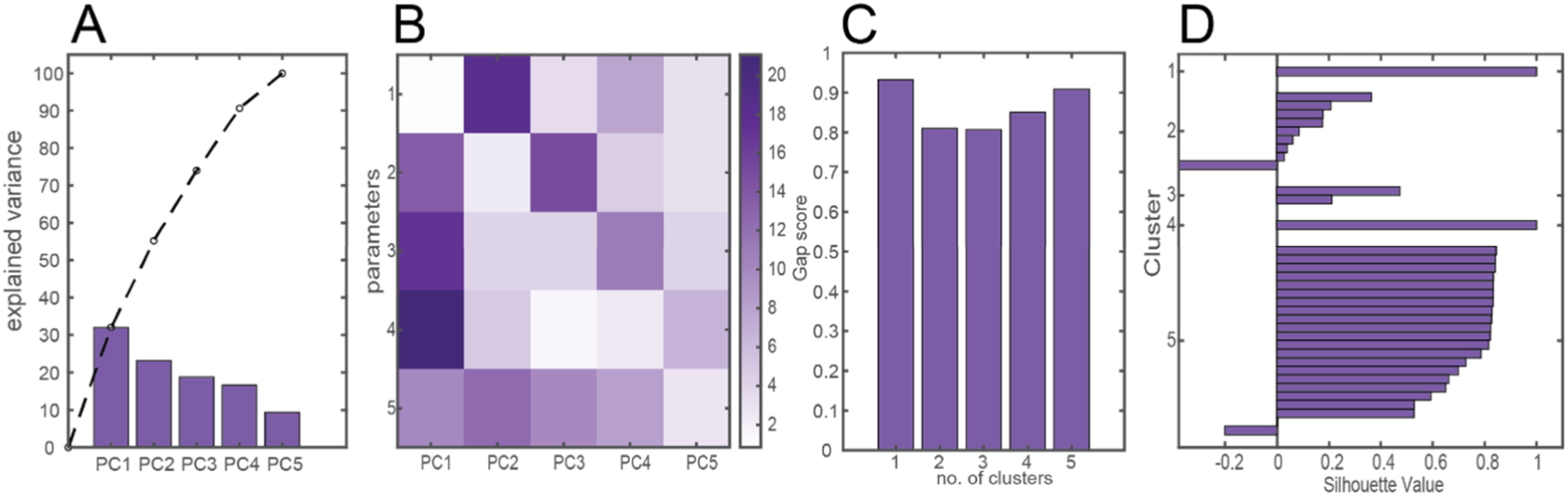
**A** Cumulative explained variance for the principal components corresponding to the spatial receptive field fit (Fig. 3). **B** Correspondence of fitting parameters (see Methods) and principal components. **C** Statistical analysis of the optimal number of clusters. **D** The silhouette plot for five clusters. Negative and low values indicate a low certainty in assigning the data points to the distinct clusters.

**Figure S3:**
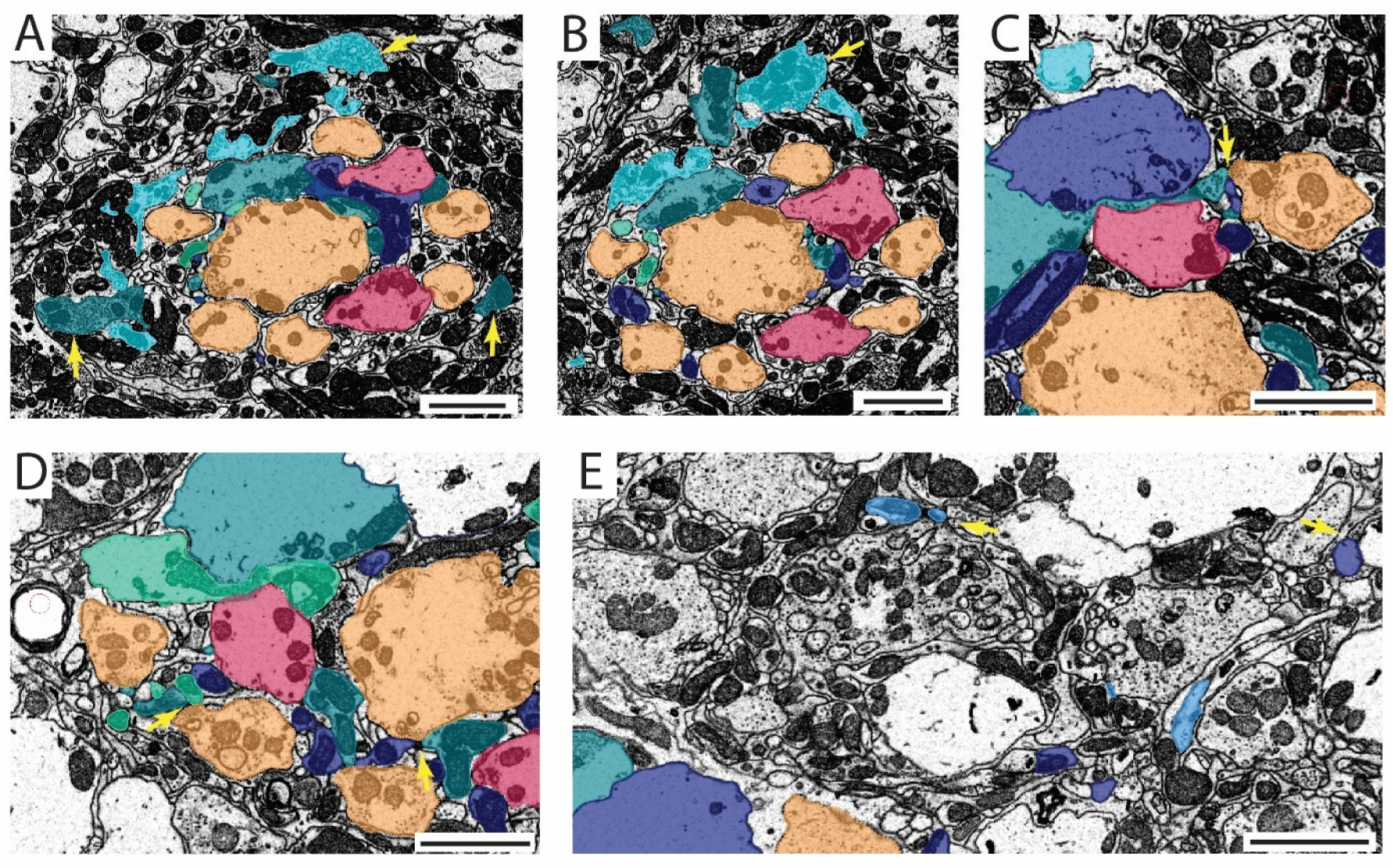
Detailed images of interesting anatomical structures in the focal cartridge. (scale bars = 2 µm). Colour code as in Fig 1 B, C. **A** vesicle filled and bulky process structures of L5 (top arrow) and L1 (two bottom arrows). **B** arrow points to vesicle filled and bulky process structures of L5. **C** in layer B: L1 (turquoise) and L2 (dark blue) are postsynaptic to a photoreceptor (orange), sharing the same synaptic site. **D** in layer B: L1 (turquoise) and L3 (green) are postsynaptic to a photoreceptor (orange). **E** L2 (dark blue) and L4 (light blue) are postsynaptic to other cells of unknown cell type outside of the focal cartridge, in the next neighbouring cartridge.

**Figure S5:**
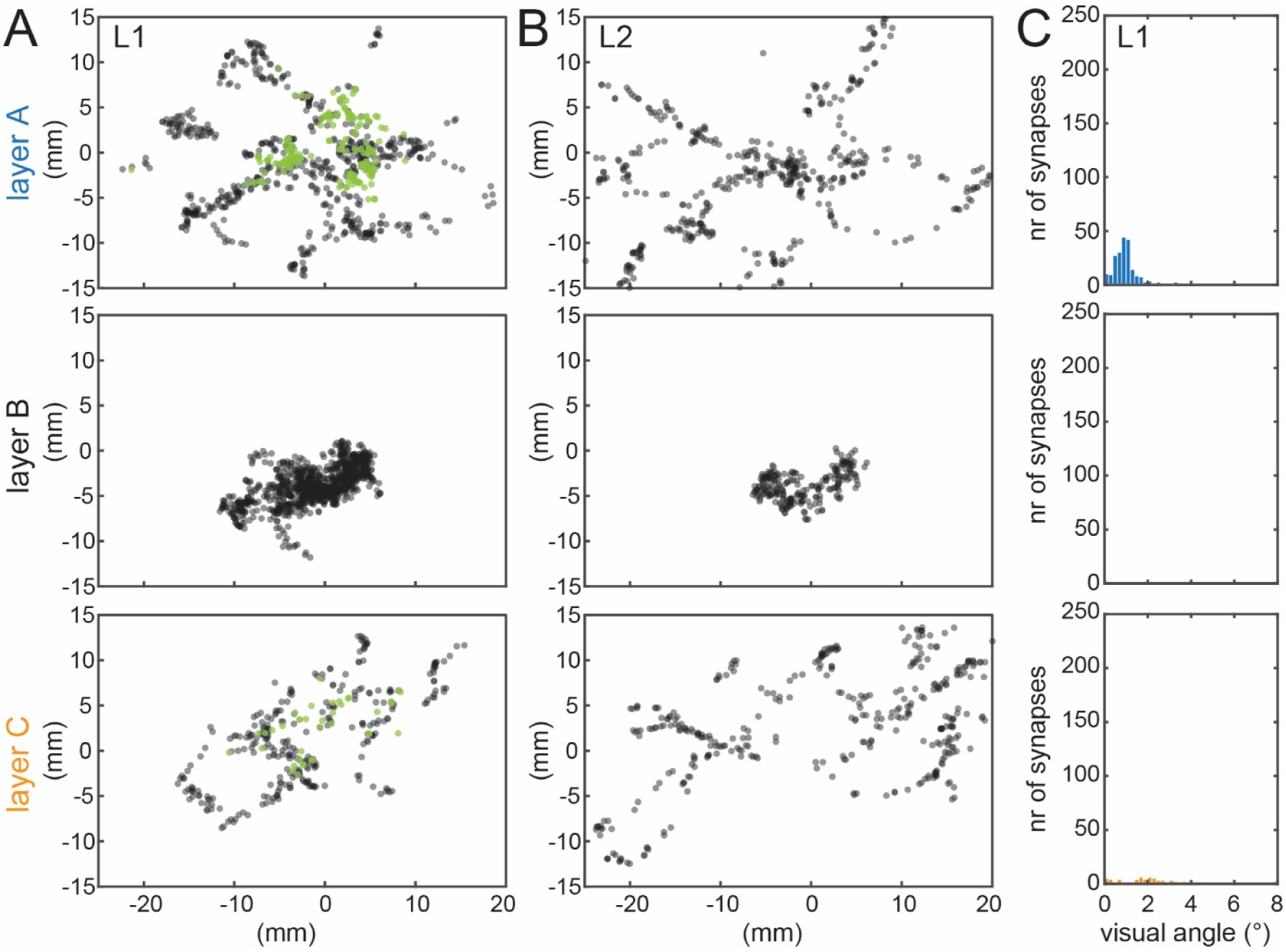
**A** Synapses of L1, in layers A, B and C (see Fig. 1C for layer definition), including pre-synapses (green) and post-synapses (black). **B** Synapses of L2. **C** Pre-synapse histogram for L1.

## Notes

### Competing Interest Statement

The authors have declared no competing interest.

### Summary of Updates

Additional analysis (computational modelling, Fig. 4), extended results and discussion section to reflect it.

## References

Behnia, R., Clark, D. A., Carter, A. G., Clandinin, T. R., & Desplan, C. (2014). Processing properties of ON and OFF pathways for Drosophila motion detection. Nature, 512(7515), 427–430.

Borst, A. (2014). Fly visual course control: behaviour, algorithms and circuits. Nat Rev Neurosci, 15(9), 590–599.

Borst, A., Haag, J., & Mauss, A. S. (2020). How fly neurons compute the direction of visual motion. J Comp Physiol A Neuroethol Sens Neural Behav Physiol, 206(2), 109–124.

Boschek, C. B. (1971). On the fine structure of the peripheral retina and lamina ganglionaris of the fly, Musca domestica. Z Zellforsch Mikrosk Anat, 118(3), 369–409.

Braitenberg, V. (1967). Patterns of projection in the visual system of the fly. I. Retina-lamina projections. Exp Brain Res, 3(3), 271–298.

Cajal, S. R. y., & Sanchez, D. (1915). Contribución al conocimiento de los centros nerviosos de los insectos.

Cardona, A., Saalfeld, S., Schindelin, J., Arganda-Carreras, I., Preibisch, S., Longair, M., … Douglas, R. J. (2012). TrakEM2 software for neural circuit reconstruction. PLoS One, 7(6), e38011.

Clark, D. A., Bursztyn, L., Horowitz, M. A., Schnitzer, M. J., & Clandinin, T. R. (2011). Defining the computational structure of the motion detector in Drosophila. Neuron, 70(6), 1165–1177.

Clark, D. A., & Demb, J. B. (2016). Parallel Computations in Insect and Mammalian Visual Motion Processing. Current Biology, 26(20), R1062–R1072.

Deerinck, T. J. (2010). NCMIR methods for 3D EM: a new protocol for preparation of biological specimens for serial block-face SEM. Microscopy [Online], 6.

Denk, W., & Horstmann, H. (2004). Serial block-face scanning electron microscopy to reconstruct three-dimensional tissue nanostructure. PLoS Biol, 2(11), e329.

Egelhaaf, M. (2023). Optic flow based spatial vision in insects. J Comp Physiol A Neuroethol Sens Neural Behav Physiol, 209(4), 541–561.

Fischbach, K.-F., & Dittrich, A. (1989). The optic lobe of Drosophila melanogaster. I. A Golgi analysis of wild-type structure. Cell and tissue research, 258(3), 441–475.

Greiner, B., Ribi, W. A., & Warrant, E. J. (2004a). Retinal and optical adaptations for nocturnal vision in the halictid bee Megalopta genalis. Cell Tissue Res, 316(3), 377–390.

Greiner, B., Ribi, W. A., & Warrant, E. J. (2005). A neural network to improve dim-light vision? Dendritic fields of first-order interneurons in the nocturnal bee Megalopta genalis. Cell Tissue Res, 322(2), 313–320.

Greiner, B., Ribi, W. A., Wcislo, W. T., & Warrant, E. J. (2004b). Neural organisation in the first optic ganglion of the nocturnal bee Megalopta genalis. Cell Tissue Res, 318(2), 429–437.

Hartline, H. K. (1969). Visual receptors and retinal interaction. Science, 164(3877), 270–278.

Joesch, M., Schnell, B., Raghu, S. V., Reiff, D. F., & Borst, A. (2010). ON and OFF pathways in Drosophila motion vision. Nature, 468(7321), 300–304.

Ketkar, M. D., Gur, B., Molina-Obando, S., Ioannidou, M., Martelli, C., & Silies, M. (2022). First-order visual interneurons distribute distinct contrast and luminance information across ON and OFF pathways to achieve stable behavior. Elife, 11.

Ketkar, M. D., Shao, S., Gjorgjieva, J., & Silies, M. (2023). Multifaceted luminance gain control beyond photoreceptors in Drosophila. Current Biology, 33(13), 2632–2645 e2636.

Kirschfeld, K. (1967). [The projection of the optical environment on the screen of the rhabdomere in the compound eye of the Musca]. Exp Brain Res, 3(3), 248–270.

Kramer, R. H., & Davenport, C. M. (2015). Lateral Inhibition in the Vertebrate Retina: The Case of the Missing Neurotransmitter. PLoS Biol, 13(12), e1002322.

Land, M. F., & Nilsson, D.-E. (2012). Animal Eyes. OUP Oxford.

Luo, L. (2021). Architectures of neuronal circuits. Science, 373(6559), eabg7285.

Macpherson, T., Matsumoto, M., Gomi, H., Morimoto, J., Uchibe, E., & Hikida, T. (2021). Parallel and hierarchical neural mechanisms for adaptive and predictive behavioral control. Neural Netw, 144, 507–521.

Matsushita, A., Stewart, F., Ilic, M., Chen, P. J., Wakita, D., Miyazaki, N., … Arikawa, K. (2022). Connectome of the lamina reveals the circuit for early color processing in the visual pathway of a butterfly. Current Biology, 32(10), 2291–2299 e2293.

Meinertzhagen, I. A., Menzel, R., & Kahle, G. (1983). The identification of spectral receptor types in the retina and lamina of the dragonflySympetrum rubicundulum. Journal of Comparative PhysiologyA, 151(3), 295–310.

Meinertzhagen, I. A., & O’Neil, S. D. (1991). Synaptic organization of columnar elements in the lamina of the wild type in Drosophila melanogaster. J Comp Neurol, 305(2), 232–263.

Mimura, K. (1976). Some spatial properties in the first optic ganglion of the fly. Journal of Comparative PhysiologyA, 105(1), 65–82.

Nassi, J. J., & Callaway, E. M. (2009). Parallel processing strategies of the primate visual system. Nat Rev Neurosci, 10(5), 360–372.

Niven, J. E., & Laughlin, S. B. (2008). Energy limitation as a selective pressure on the evolution of sensory systems. J Exp Biol, 211(Pt 11), 1792–1804.

Ohly, K. P. (1975). The neurons of the first synaptic region of the optic neuropil of the firefly, Phausis splendidula l. (Coleoptera). Cell Tissue Res, 158(1), 89–109.

Priebe, N. J., & Ferster, D. (2008). Inhibition, spike threshold, and stimulus selectivity in primary visual cortex. Neuron, 57(4), 482–497.

Pyza, E., & Meinertzhagen, I. A. (1995). Monopolar Cell Axons in the First Optic Neuropil of the Housefly, Musca-Domestica L, Undergo Daily Fluctuations in Diameter That Have a Circadian Basis. Journal of Neuroscience, 15(1), 407–418.

Ribi, W. A. (1975a). The first optic ganglion of the bee. Cell and tissue research, 165(1), 103–111.

Ribi, W. A. (1975b). Golgi studies of the first optic ganglion of the ant, Cataglyphis bicolor. Cell Tissue Res, 160(2), 207–217.

Ribi, W. A. (1977). Fine structure of the first optic ganglion (lamina) of the cockroach, Periplaneta americana. Tissue Cell, 9(1), 57–72.

Ribi, W. A. (1978). Gap junctions coupling photoreceptor axons in the first optic ganglion of the fly. Cell Tissue Res, 195(2), 299–308.

Ribi, W. A. (1987). Anatomical Identification of Spectral Receptor Types in the Retina and Lamina of the Australian Orchard Butterfly, Papilio-Aegeus-Aegeus D. Cell and tissue research, 247(2), 393–407.

Rivera-Alba, M., Vitaladevuni, S. N., Mishchenko, Y., Lu, Z., Takemura, S. Y., Scheffer, L., … de Polavieja, G. G. (2011). Wiring economy and volume exclusion determine neuronal placement in the Drosophila brain. Current Biology, 21(23), 2000–2005.

Ryu, L., Kim, S. Y., & Kim, A. J. (2022). From Photons to Behaviors: Neural Implementations of Visual Behaviors in Drosophila. Front Neurosci, 16, 883640.

Schindelin, J., Arganda-Carreras, I., Frise, E., Kaynig, V., Longair, M., Pietzsch, T., … Cardona, A. (2012). Fiji: an open-source platform for biological-image analysis. Nat Methods, 9(7), 676–682.

Shimohigashi, M., & Tominaga, Y. (1999). Synaptic organization in the lamina of the superposition eye of a skipper butterfly, Parnara guttata. J Comp Neurol, 408(1), 107–124.

Stöckl, A. L., O’Carroll, D., & Warrant, E. J. (2017). Higher-order neural processing tunes motion neurons to visual ecology in three species of hawkmoths. Proc Biol Sci, 284(1857), 20170880.

Stöckl, A. L., O’Carroll, D. C., & Warrant, E. J. (2016a). Neural Summation in the Hawkmoth Visual System Extends the Limits of Vision in Dim Light. Current Biology, 26(6), 821–826.

Stöckl, A. L., O’Carroll, D. C., & Warrant, E. J. (2020). Hawkmoth lamina monopolar cells act as dynamic spatial filters to optimize vision at different light levels. Sci Adv, 6(16), eaaz8645.

Stöckl, A. L., Ribi, W. A., & Warrant, E. J. (2016b). Adaptations for nocturnal and diurnal vision in the hawkmoth lamina. J Comp Neurol, 524(1), 160–175.

Strausfeld, N. J. (1970). Golgi studies on insects Part II. The optic lobes of Diptera. Philosophical Transactions of the Royal Society of London. B, Biological Sciences, 258(820), 135–223.

Strausfeld, N. J. (2012). Arthropod brains: evolution, functional elegance, and historical significance. Belknap Press.

Strausfeld, N. J., & Blest, A. (1970). Golgi studies on insects Part I. The optic lobes of Lepidoptera. Philosophical Transactions of the Royal Society of London. B, Biological Sciences, 258(820), 81–134.

Strausfeld, N. J., & Braitenberg, V. (1970). The compound eye of the fly (Musca domestica): connections between the cartridges of the lamina ganglionaris. Zeitschrift für vergleichende Physiologie, 70(2), 95–104.

Strausfeld, N. J., & Campos-Ortega, J. A. (1977). Vision in insects: pathways possibly underlying neural adaptation and lateral inhibition. Science, 195(4281), 894–897.

Wakita, D., Shibasaki, H., Kinoshita, M., & Arikawa, K. (2024). Morphology and spectral sensitivity of long visual fibers and lamina monopolar cells in the butterfly Papilio xuthus. J Comp Neurol, 532(2), e25579.

Warrant, E. J. (1999). Seeing better at night: life style, eye design and the optimum strategy of spatial and temporal summation. Vision Res, 39(9), 1611–1630.

Watanabe, M., & Rodieck, R. W. (1989). Parasol and midget ganglion cells of the primate retina. J Comp Neurol, 289(3), 434–454.

Wolburg-Buchholz, K. (1979). The organization of the lamina ganglionaris of the hemipteran insects, Notonecta glauca, Corixa punctata and Gerris lacustris. Cell Tissue Res, 197(1), 39–59.

Zettler, F., & Jarvilehto, M. (1972). Lateral Inhibition in an Insect Eye. Zeitschrift für vergleichende Physiologie, 76(3), 233-+.

